# Could radical pairs play a role in xenon-induced general anesthesia?

**DOI:** 10.1101/2020.08.24.265082

**Authors:** Jordan Smith, Hadi Zadeh Haghighi, Christoph Simon

## Abstract

Understanding the mechanisms underlying anesthesia would be a key step towards understanding consciousness. The process of xenon-induced general anesthesia has been shown to involve electron transfer, and the potency of xenon as a general anesthetic exhibits isotopic dependence. We propose that these observations can be explained by a mechanism in which the xenon nuclear spin influences the recombination dynamics of a naturally occurring radical pair of electrons. We develop a simple model inspired by the body of work on the radical-pair mechanism in cryptochrome in the context of avian magnetoreception, and we show that our model can reproduce the observed isotopic dependence of the general anesthetic potency of xenon in mice. Our results are consistent with the idea that radical pairs of electrons with entangled spins could be important for consciousness.

## 1 Introduction

Understanding consciousness remains one of the big open questions in neuroscience^1^, and in science in general. The study of anesthesia is one of the key approaches to elucidating the processes underlying consciousness^2,3^, but there are still significant open questions regarding the physical mechanisms of anesthesia itself^4^.

One anesthetic agent that has been studied extensively is xenon. Xenon has been shown experimentally to produce a state of general anesthesia in several species, including *Drosophila^5^,* mice^6^, and humans^7^. While the anesthetic properties of xenon were discovered in 1939^8^, the exact underlying mechanism by which it produces anesthetic effects remains unclear even after decades of research^9^. Our focus here is on this underlying physical mechanism, and there are important hints about the mechanism provided by two recent publications.

First, Turin *et al.* showed that when xenon acts anesthetically on *Drosophila* unpaired electron spin resonance takes place^5^, providing evidence of some form of electron transfer. Turin *et al.* proposed that the anesthetic action of xenon may be caused by the xenon atom(s) acting as an “electron bridge“^5^, facilitating the electron transfer between a nearby electron donor and electron acceptor. They supported their proposal by density-functional theory calculations showing the effect of xenon on nearby molecular orbitals. Second, Li *et al.* showed experimentally that isotopes of xenon with non-zero nuclear spin had reduced anesthetic potency in mice compared with isotopes with no nuclear spin^6^.

If the process by which xenon produces anesthetic effects includes free-electron transfer as well as nuclear-spin dependence, a mechanistic framework proposed to explain xenon-induced anesthesia should possess these characteristics. Here we show that a model involving a radical pair of electrons (RP) and the subsequent modulation of the RP spin dynamics by hyperfine interactions is consistent with these assumptions.

The radical pair mechanism (RPM) was first proposed more than 50 years ago^10^. The rupture of a chemical bond can create a pair of electrons that are localized on two different molecular fragments, but whose spins are entangled in a singlet state^11^. The magnetic dipole moment associated with electron spin can interact and couple with other magnetic dipoles, including hyperfine interactions and Zeeman interactions^12^. As a consequence of such interactions, the initial singlet state can evolve into a more complex state that has both singlet and triplet components. Eventually, the coherent oscillation of the RP between singlet and triplet states ceases and the electrons may recombine (for the singlet component of the state) or diffuse apart to form various triplet products^13^. In order for significant interconversion between singlet and triplet states to occur, the RP lifetime and the RP spin-coherence lifetime should be comparable to the electron Larmor precession period, which in the case of the geomagnetic field of the Earth (*B* ≈ 50*μ*T) is approximately 700 ns^13–15^. The coherent spin dynamics and spin-dependent reactivity of radical pairs allow magnetic interactions which are 6 orders of magnitude smaller than the thermal energy, *k_B_T*, to have predictable and reproducible effects on chemical reaction yields^14,16^.

The RPM has become a prominent concept in quantum biology^17^. In particular, it has been studied in detail for the cryptochrome protein as a potential explanation for avian magnetoreception^16,18–23^. In the present work we apply the principles and methods used to investigate cryptochrome^14^ to xenon-induced anesthesia.

General anesthetics produce widespread neurodepression in the central nervous system by enhancing inhibitory neurotransmission and reducing excitatory neurotransmission across synapses^24,25^. Three ligand-gated ion-channels in particular have emerged as likely molecular targets for a range of anesthetic agents^26^: the inhibitory glycine receptor, the inhibitory *γ*-aminobutyric acid type-A (GABAA) receptor, and the excitatory N-methyl-D-aspartate (NMDA) receptor^27^. NMDA receptors require both glycine and glutamate for excitatory activation^28^, and it has been suggested that xenon’s anesthetic action is related to xenon atoms participating in competitive inhibition of the NMDA receptor in central-nervous-system neurons by binding at the glycine binding site in the NMDA receptor^26,27,29,30^. Armstrong *et al.* propose that the glycine binding site of the NMDA receptor contains aromatic rings^26,29^ of similar chemical nature to those found in cryptochrome, which is consistent with the possibility that the anesthetic action of xenon uses a spin-dependent process similar to the radical-pair mechanism (RPM) thought to occur in cryptochrome. Let us note that other targets have also been proposed for many anesthetics, e.g. tubulin^31^. In the following we focus on the NMDA receptor to be specific, but our model is very general and could well apply to other targets.

We suggest that the electron transfer evidenced by Turin et al.^5^ could affect the recombination dynamics of a naturally occurring radical pair (based on the results of Armstrong et *al.* this RP may involve aromatic phenylalanine residues^29^ located in the binding site), and that an isotope of xenon with a non-zero nuclear spin could couple with the electron spins of such a radical pair, reducing the “electron bridge” effect that is proposed to correlate with anesthesia. Such a mechanism is consistent with the experimental results of Li *et al*.^6^ that xenon isotopes with non-zero nuclear spin have reduced anesthetic potency compared to isotopes with zero nuclear spin.

We hypothesize that in the context of xenon-induced general anesthesia, xenon itself may not be involved in the creation of radical pairs, where the energy for radical-pair creation likely comes from another source such as local ROS. It has been suggested that water (a source of oxygen) may be present in the NMDA receptor^26,28^, and Aizenman et al.^32^ as well as Girouard et al.^33^ suggest that reactive oxygen species (ROS) may be located in the NMDA receptor. Further, Turin and Skoulakis found that when a sample of xenon gas was administered to *Drosophila* without oxygen gas present in the sample no spin changes were observed in the flies^34^. Here we propose that ROS could be involved in the formation and existence of a naturally occurring RP in the NMDA receptor.

Given that cryptochrome is one case in which the RPM has been studied extensively, we propose that by recognizing commonalities in the biological and chemical environments in which magnetoreception and xenon-induced general anesthesia are thought to take place, analytical and numerical techniques that have been used to study cryptochrome may be adapted and applied to the case of xenon-induced anesthesia, potentially providing insight into general anesthetic mechanisms. We explore the feasibility of such a mechanism by determining and analyzing the necessary parameters and conditions under which the spin-dependent RP product yields can explain the experimental isotope-dependent anesthetic effects reported by Li *et al^6^*.

## 2 Results

### Predicting Experimental Xenon Anesthesia Results using the RPM Model

#### Quantifying anesthetic potency

In the work of Li et al.^6^, a metric referred to as the “loss of righting reflex ED50” (LRR-ED50) was defined using the concentration of xenon administered to mice, in which the mice were no longer able to right themselves within 10 s of being flipped onto their backs. The LRR-ED50 metric was reported to be correlated with consciousness in mice, and was measured experimentally for ^132^Xe, ^134^Xe, ^131^Xe, and ^129^Xe to be 70(4)%, 72(5)%, 99(5)%, and 105(7)%, respectively^6^.

Here we defined the anesthetic potency as the inverse of the LRR-ED50 metric. In order to quantify the anesthetic potency of the various xenon isotopes, the potency of ^132^Xe (with *I =* 0) was normalized to 1. The inverse of the LRR-ED50 value of isotopes ^131^Xe and ^129^Xe were then divided by that of ^132^Xe. The relative isotopic anesthetic potencies were quantified as *Pot*_0_ = 1, *Pot*_3/2_ = 0.71(8), and *Pot*_1/2_ = 0.67(8) for ^132^Xe, ^131^Xe, and ^129^Xe, respectively, as shown in Table 1, where *Pot*_0_ is the relative potency of xenon with nuclear spin *I* = 0, and likewise for *Pot*_3/2_ and *Pot*_1/2_.

**Table 1.** Xenon isotopic nuclear spin, LRR-ED50, and Pot values for xenon isotopes ^132^Xe, ^134^Xe, ^131^Xe, and ^129^Xe. LRR-ED50 values as reported in the work of Li *et al*.^6^

| Isotope | Nuclear Spin, $I$ | LRR-ED50 (%) | $Pot$ |
| --- | --- | --- | --- |
| $^{132}\text{Xe}$ | 0 | 70(4) | 1 |
| $^{134}\text{Xe}$ | 0 | 72(5) | - |
| $^{131}\text{Xe}$ | 3/2 | 99(5) | 0.71(8) |
| $^{129}\text{Xe}$ | 1/2 | 105(7) | 0.67(8) |

#### RPM model

We have developed an RPM model to predict anesthetic potency by making a connection with the relative singlet yield for different isotopes, as described in more detail below. The model that we have used here was developed using a hypothetical xenon-NMDA receptor RP system based on the information about xenon action sites mentioned previously, involving xenon atoms surrounded by phenylalanine and tryptophan residues located in the glycine-binding site of the NMDA receptor, and also modelled after the cryptochrome case as related to magnetoreception. A spin-correlated radical pair of electrons (termed electrons A and B), most likely found in chemically excited phenylalanine residues^26^, couple to xenon nuclei with non-zero nuclear spin, as shown in Fig. 1.

**Figure 1.**
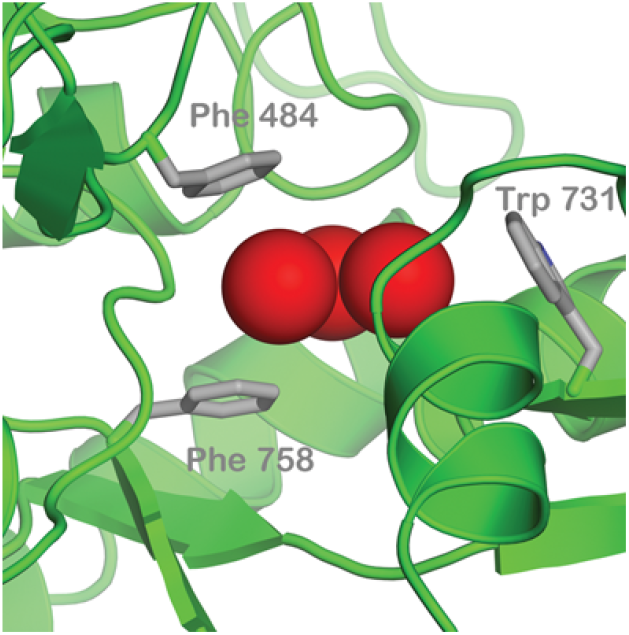
Aromatic residues are important for the binding of xenon and glycine at the glycine binding site of the NMDA receptor. The image shows the predicted position of xenon atoms (red spheres) in the glycine site together with the aromatic residues phenylalanine 758, phenylalanine 484, and tryptophan 731.^29^

The number of xenon atoms located in the active site when anesthetic action takes place is not yet completely clear, and the work of Dickinson *et al.* suggests that the number of xenon atoms simultaneously present in the active site ranges probabilistically between zero and three^26^.

Here we show that the simplest case of a single xenon atom occupying the active site already allows us to explain the anesthetic potency ratios derived from the experimental results of Li *et al.*^6^. The additional degrees of freedom implicit in more complex Hilbert spaces, such as the cases of two and three-xenon occupation states, only aid the model in explaining the experimental results of Li *et al.*. Further, when our model is optimized in order to reproduce the experimental results, the two-xenon occupation state essentially reduces to the single-xenon occupation state, in which one hyperfine interaction between a xenon nucleus and one of the radical electrons is dominant. We therefore focus our analysis on the single atom case, but the two and three-xenon occupation states are discussed in the Supplementary Information.

The Hamiltonian of the RPM in the case of xenon-induced anesthesia depends not only upon the number of xenon atoms present in the glycine binding site of the NMDA receptor, but also upon the assumed hyperfine interactions. In the simulation involving a single xenon atom occupying the active site it was assumed that the xenon nucleus may couple to both radical electrons, and the Hamiltonian is given as

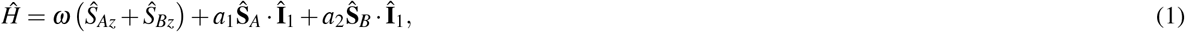

where 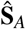 and 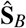 are the spin operators of radical electrons A and B, respectively, 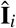 is the nuclear spin operator of xenon nucleus *i, a_j_* is a hyperfine coupling constant where 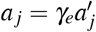, and *ω* is the Larmor precession frequency of the electrons about an external magnetic field^14^. The Larmor precession frequency is defined as *ω = γ_e_B*, where *γ_e_* is the gyromagnetic ratio of an electron and B is the external magnetic field strength. It should be pointed out that we focus here on the hyperfine interactions between the radical electrons and xenon atoms, and that interactions between the two electron spins, as well as potential interactions between the electron spins and other nuclei are neglected. This is justifiable for our purposes, since we are primarily interested in the differential effect of the xenon nuclear spin.

#### Determination of Singlet Yield Ratios

The eigenvalues and eigenvectors of the Hamiltonian can be used to determine the ultimate singlet yield (Φ_*S*_) for all times much greater than the radical-pair lifetime (*t* ≫ *τ*):^14^

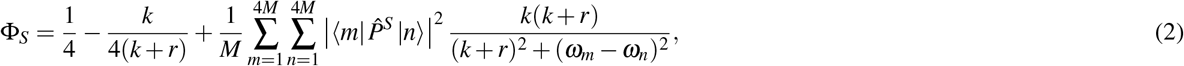

where, following the methodology used by Hore^14^ in the context of cryptochrome, *M* is the total number of nuclear spin configurations, 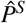 is the singlet projection operator, |*m*〉 and |*n*〉 are eigenstates of 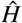 with corresponding energies of 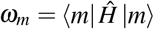 and 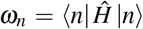, respectively, *k = τ*^−1^ is inverse of the RP lifetime, and 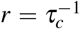 is the inverse of the RP spin-coherence lifetime.

The spin-dependent RP singlet yield was calculated for each xenon isotope under consideration. The singlet yield ratio (*SR*) for each nuclear spin value was then calculated by dividing the singlet yield obtained using the given spin value by the singlet yield obtained using spin *I* = 0, resulting in the ratio of spin-1/2 singlet yield to spin-0 singlet yield being expressed as *SR*_1/2_, and the ratio of spin-3/2 singlet yield to spin-0 singlet yield expressed as *SR*_3/2_, with the singlet yield ratio of spin-0 being normalized to *SR*_0_ = 1. The calculated singlet yield ratios were compared with the xenon potency ratios (*Pot*) derived using the data reported by Li *et al*.^6^ as described above.

We investigated the sensitivity of the singlet yield ratios to changes in the hyperfine interactions (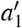 and 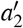), RP reaction rate (*k*), external magnetic field strength (*B*), and RP spin-coherence relaxation rate (*r*). The dependence of the quantity |*Pot − SR*| on 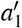 and 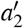 is shown in Figs. 2/3, while the dependence of |*Pot − SR*| on the relationship between *r* and *k* is shown in Figs. 4/5. The relationship between *SR* and *B* can be seen in Fig. 6. The same dependencies for the two-xenon occupation state can be seen in Supplementary Figs. S1/S2, S3/S4, and S5, respectively. Our goal was to find regions in parameter space such that the spin-dependent singlet yield ratios match the anesthetic isotopic potency ratios, i.e., the quantities |*Pot*_1/2_ − *SR*_1/2_| and |*Pot*_3/2_ − *SR*_3/2_| should be smaller than the experimental uncertainties on the anesthetic potency.

**Figure 2.**
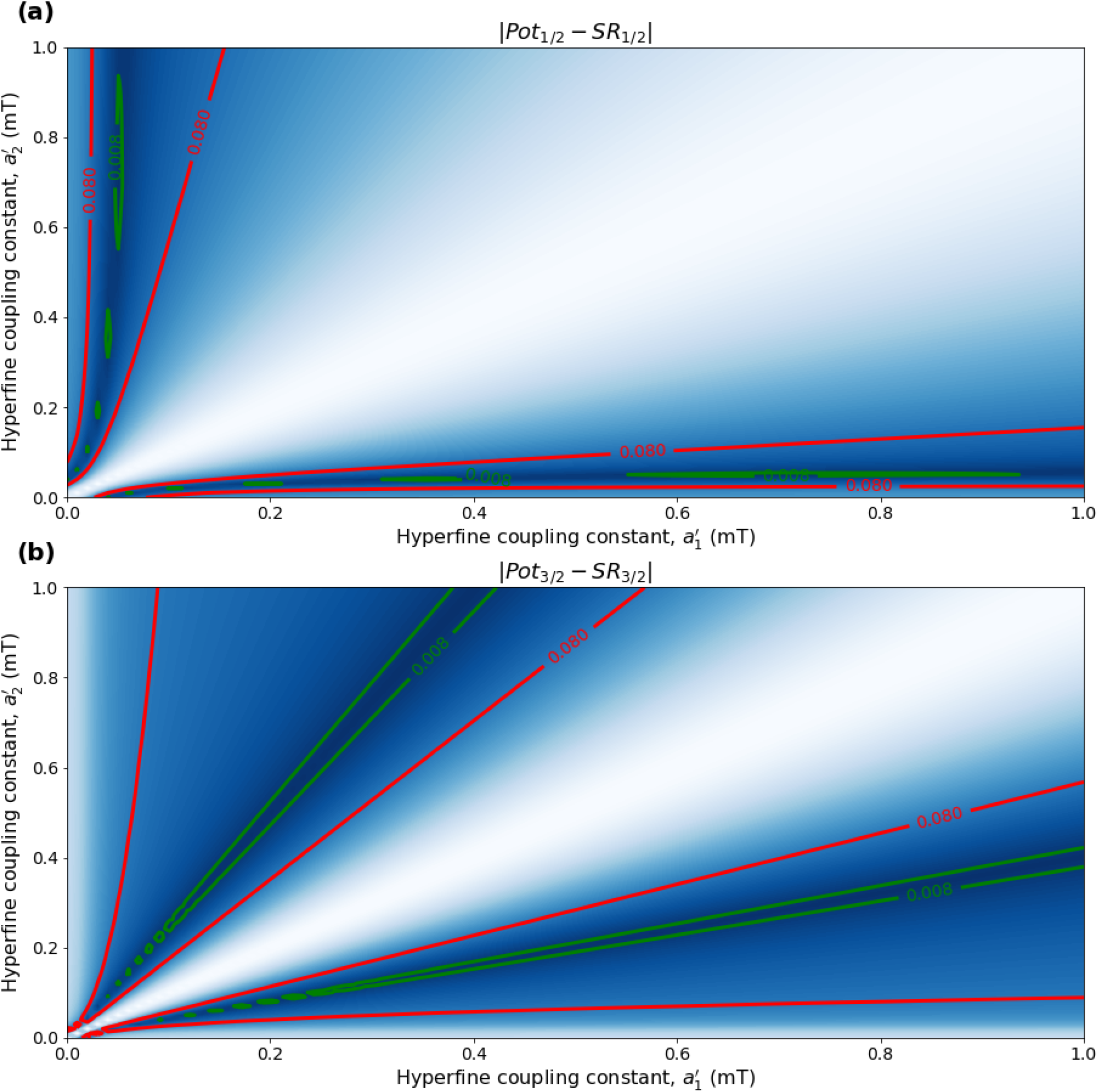
The dependence of the single-xenon RPM model on changes in the hyperfine coupling constants 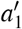 and 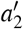 for 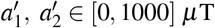, using *r* = 1.0 × 10^6^ s^−1^, *B* = 50 *μ*T, and *τ* = 1.0 × 10^−6^ s. The model can explain the experimentally derived^6^ relative anesthetic potency of xenon for values of *r* and *k* where |*Pot*_1/2_ − *SR*_1/2_|, |*Pot*_3/2_ − *SR*_3/2_| ≤ 0.08. **(a)** The absolute difference between *Pot*_1/2_ and *SR*_1/2_. **(b)** The absolute difference between *Pot*_3/2_ and *SR*_3/2_.

**Figure 3.**
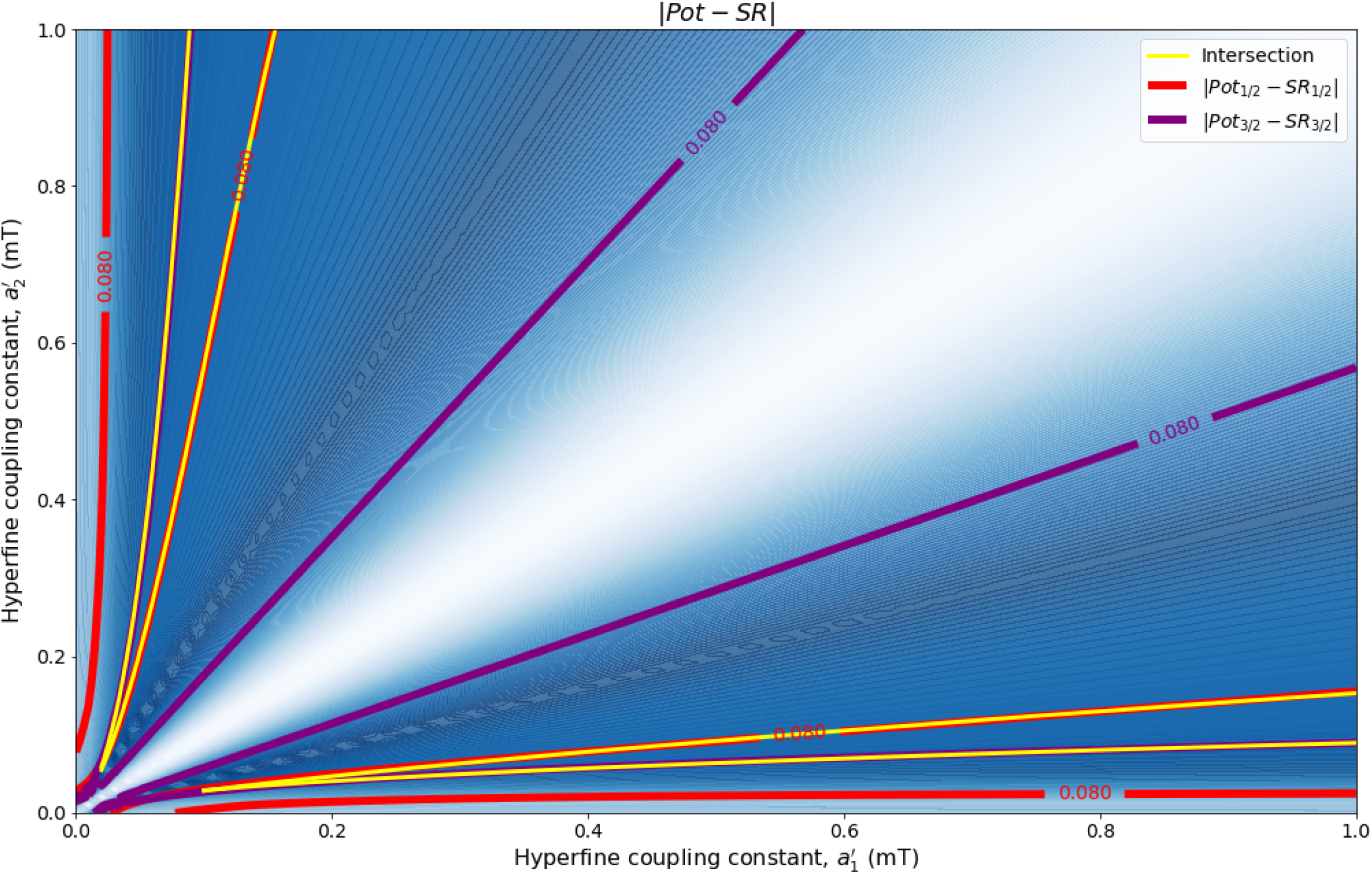
The dependence of the single-xenon RPM model on changes in the hyperfine coupling constants 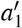 and 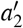 for 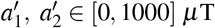, using *r* = 1.0 × 10^6^ s^−1^, *B* = 50 *μ*T, and τ = 1.0 × 10^−6^ s. The model can explain the experimentally derived^6^ relative anesthetic potency of xenon where |*Pot*_1/2_ − *SR*_1/2_| ≤ 0.080 and |*Pot*_3/2_ − *SR*_3/2_| ≤ 0.08 intersect.

**Figure 4.**
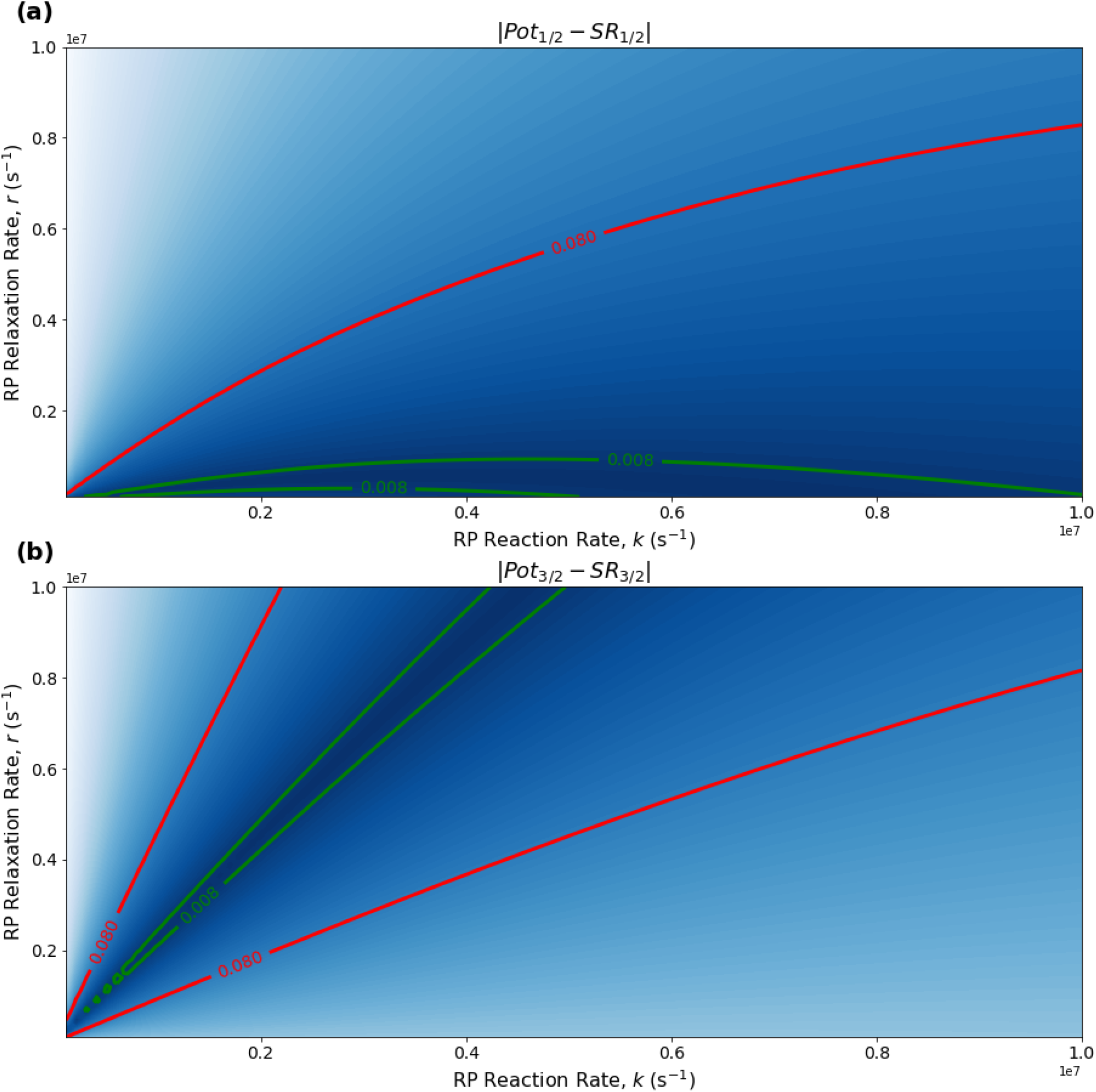
The dependence of the single-xenon RPM model on the relationship between *r* and *k* for *r, k* ∈ [1.0 × 10^5^,1.0 × 10^7^] s^−1^, using 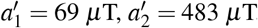, and *B* = 50 *μ*T. The model can explain the experimentally derived^6^ relative anesthetic potency of xenon for values of *r* and *k* where |*Pot*_1/2_ − *SR*_1/2_|, |*Pot*_3/2_ − *SR*_3/2_| ≤ 0.08. **(a)** The absolute difference between *Pot*_1/2_ and *SR*_1/2_. **(b)** The absolute difference between *Pot*_3/2_ and *SR*_3/2_.

**Figure 5.**
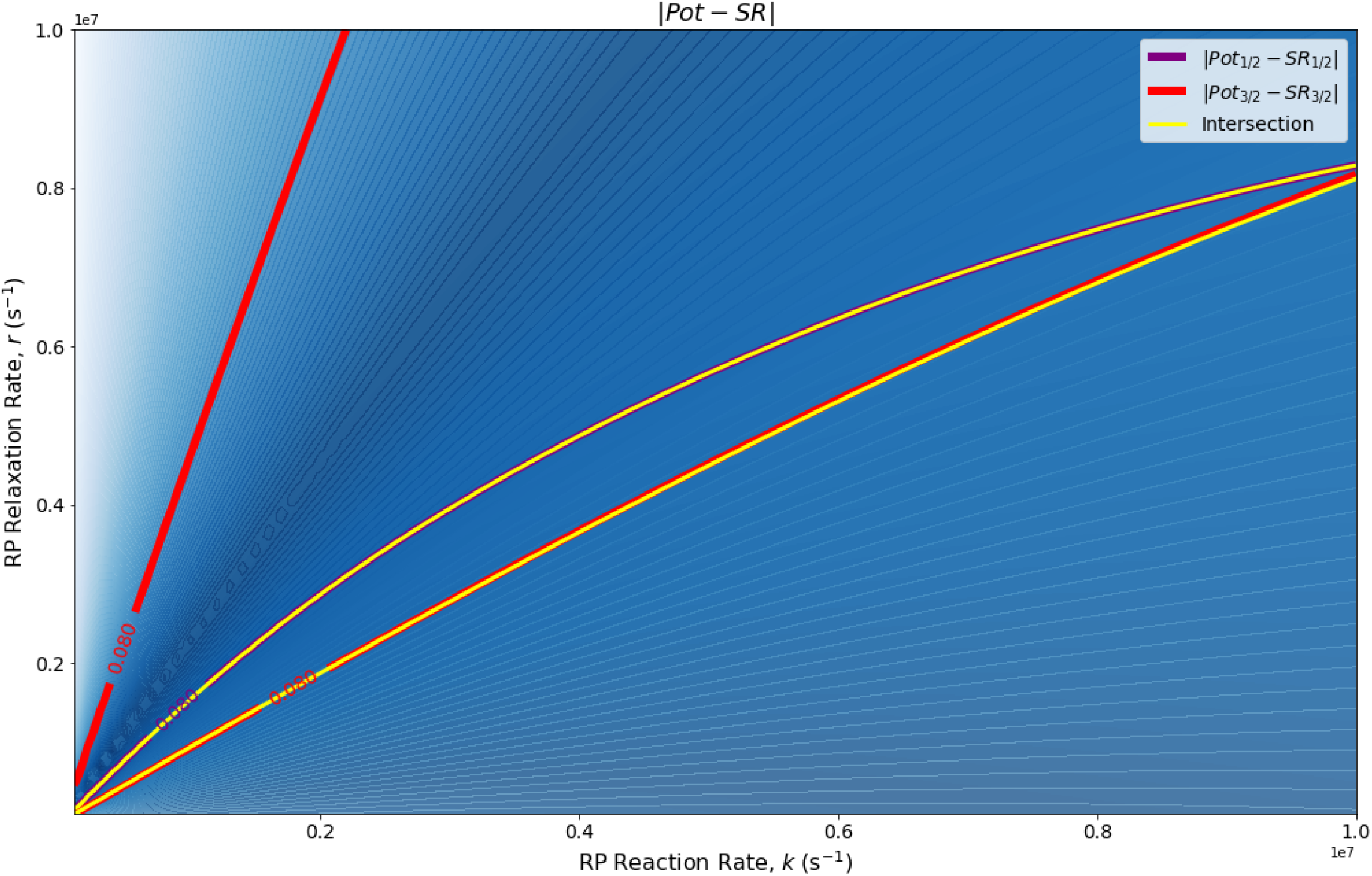
The dependence of the single-xenon RPM model on the relationship between *r* and *k* for r, *k* ∈ [1.0 × 10^5^, 1.0 × 10^7^] s^−1^, using 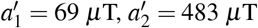, and *B* = 50 *μ*T. The model can explain the experimentally derived^6^ relative anesthetic potency of xenon where |*Pot*_1/2_ − *SR*_1/2_| ≤ 0.080 and |*Pot*_3/2_ − *SR*_3/2_| ≤ 0.08 intersect.

**Figure 6.**
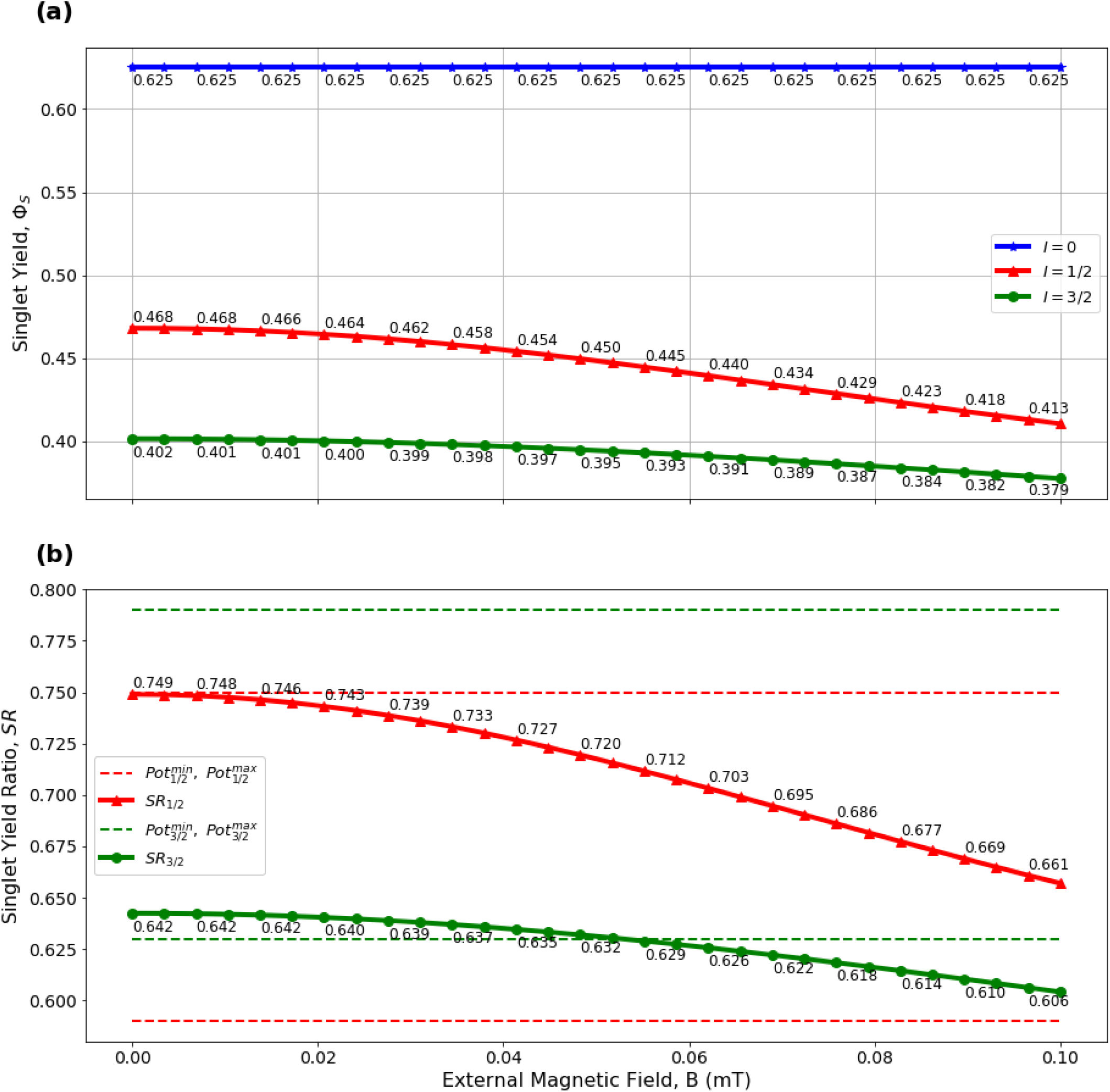
The external field-dependence of the single-xenon RPM model with *B* ∈ [0, 100] *μ*T, *τ* = 1.0 × 10^−6^ s, *r* = 1.0 × 10^6^ s^−1^, and hyperfine constants of 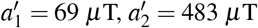. **(a)** The absolute singlet yield using xenon isotopic nuclear spin values of *I* ∈ {0, 1/2, 3/2}. **(b)** The relative singlet yield ratios *SR*_1/2_ and *SR*_3/2_ with the singlet yield of spin *I* = 0 being normalized to *SR*_0_ = 1. Values of *SR* and *Pot* agree for *B* ∈ [0, 51] *μ*T.

In the case of single-xenon occupation the optimized parameter values were found to be 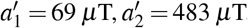, *B* = 50 *μ*T, *k* = *r* = 1.0 × 10^6^ s^−1^, resulting in *SR*_1/2_ = 0.72 and *SR*_3/2_ = 0.63. It should be noted here that while |*Pot*_3/2_ − *SR*_1/2_| is greater than |*Pot*_3/2_ − *SR*_1/2_| for these optimized parameter values, both *SR*_3/2_ and *SR*_1/2_ are consistent with the experimental results of Li *et al*.^6^, within their uncertainties. Further experiments with smaller uncertainty about the relative anesthetic potencies of spin-3/2 and spin-1/2 xenon isotopes would be of interest.

## 3 Discussion

The optimized parameter values for a single xenon atom suggest strong coupling only between the xenon nucleus and one of the two RP electrons, while having weak interactions with the other, see Figs. 2 and 3. To gain a deeper understanding of this coupling, it would be of interest to perform molecular modelling of the electron transfer between xenon and tryptophan/phenylalanine residues using Marcus Theory. Such modelling could be expanded on by exploring the molecular dynamics of the binding site using quantum mechanics/molecular mechanics (QM/MM) simulation techniques, similar to those used in the case of cryptochrome^35^, to determine which aromatic rings are most likely to contain a radical pair, and therefore which molecules the xenon nuclei are most likely to couple with. Further, rather than determining the hyperfine coupling constants by fitting parameters to a model, it could be useful to perform density functional theory (DFT) modelling to determine theoretical hyperfine coupling constants, accounting for the quantity and relative positioning of the xenon atoms within the NMDA receptor glycine binding site.

The sensitivity analysis of the relationship between the RP spin-coherence relaxation rate (r) and the RP first-order reaction rate (*k*) shown in Figs. 4 and 5 is promising, in that the RP lifetime requirements are within an order of magnitude of the results reported by Hore in the cryptochrome case^14^. The calculated range for the RP lifetime, *τ*, is comparable to the spin-coherence lifetime, *τ_c_*, and also to the electron Larmor precession period. It is interesting to note that the *r* and *k* parameter spaces for which *SR* and *Pot* agree indicate that for spin *I* = 1/2 the RP reaction rate (*k*) should be at least as fast as the RP spin-coherence relaxation rate (*r*), while in the spin *I* = 3/2 case the inverse relation appears to be true. For both RP spin-coherence relaxation rates and RP reaction rates much above or below the value outlined by Hore^14^ of *r* = *k* = 1.0 × 10^6^ s^−1^, *SR* and *Pot* diverge. Relaxation rates much greater than these values result in the RP not having sufficient time to coherently oscillate between singlet and triplet states such that the singlet yield ratios and the derived anesthetic potency ratios are in agreement. As with the cryptochrome case, as the RP lifetime becomes much greater than the RP spin-coherence lifetime, the RP exponentially decays toward the equilibrium state of Φ_*S*_ = 0.25 for all nuclear spin values. These results emphasizes the importance of the RP spin-coherence lifetime (*τ_c_*) being comparable to the electron Larmor precession period (*τ_L_* = 2*π/ω*) and also to the requisite RP lifetime (*τ*).

When considering the sensitivity of the single-xenon model to changes in external magnetic field as shown in Fig. 6, the range of *B* values that produce agreement between *SR* and Pot ratios is given as *B* ∈ [0,51] *μ*T, and approximates the geomagnetic field at different geographic locations (25 to 65 *μ*T)^36^. This result indicates that for a single-xenon occupation state external field values vastly stronger than the geomagnetic field may result in the anesthetic potency of xenon being reduced significantly. Further, at geographic locations on the Earth where the geomagnetic field is larger than 51 *μ*T, anesthetic potency may be reduced compared to locations where *B_earth_* ≤ 51 *μ*T. It is also seen in Fig. 6 that the isotope of xenon with spin *I* = 0 has no external magnetic field dependence, while isotopes with non-zero spin do show a dependence on *B*. It would be of interest to investigate the experimental effects of the external magnetic field strength on xenon-induced general anesthesia *in vivo*. For example, such an experiment could involve the measurement of anesthetic potency of xenon isotopes with various nuclear spin values, including both zero spin and non-zero spin, in an environment with controllable external magnetic field.

In this study the RPM model in the context of cryptochrome^14^ was adapted to the case of xenon in the glycine binding site of the N-methyl-D-aspartate receptor, and the experimental isotope-dependent anesthetic potency ratios derived from the results of Li *et al*.^6^ were reproduced theoretically using the RPM model in which a single xenon atom occupies the active site of the NMDA receptor. The RP lifetime and hyperfine coupling parameters were optimized to reproduce the experimental results of Li *et al*., and the sensitivity of the model to changes in hyperfine coupling constants, RP reaction rate, external magnetic field strength, and RP spin-coherence relaxation rate was explored. The optimized model parameter values and parameter spaces found here seem physiologically feasible, and indicate that the RPM may provide a reasonable explanation for the general anesthetic action of xenon *in vivo*.

Our results thus suggest that xenon-induced general anesthesia may fall within the realm of quantum biology, and be similar in nature to the proposed mechanism of magnetoreception involving the cryptochrome protein^18^.

This also raises the question whether other anesthetic agents use the same mechanism as xenon to induce general anesthesia. It could prove interesting to explore isotopic nuclear-spin effects as well as magnetic field effects in experiments with other general anesthetic agents that are thought to function similarly to xenon, such as nitrous oxide and ketamine^37,38^.

General anesthesia is clearly related to consciousness, and it has been proposed that consciousness (and other aspects of cognition) could be related to large-scale entanglement^39–42^. Radical pairs are entangled and could be a key element in the creation of such large-scale entanglement, especially when combined with the suggested ability of axons to serve as waveguides for photons^43^. Viewed in this - admittedly highly speculative - context, the results of the present study are consistent with the idea that general anesthetic agents, such as xenon, could interfere with this large-scale entanglement process, and thus with consciousness.

## Data Availability

The datasets generated and analysed during the current study are available from the corresponding author on reasonable request.

## Acknowledgements

The authors would like to thank Peter Hore, Robert Dickinson, Dennis Salahub, Sourabh Kumar, Parisa Zarkeshian, Sumit Goswami, Faezeh Kimiaee Asadi, Stephen Wein, Jiawei Ji, Yufeng Wu, and Kenneth Sharman for their input, comments, and insights on this topic. C.S. particularly thanks Stuart Hameroff for bringing the results of Li et al.^6^ to his attention. This work was supported by the Natural Sciences and Engineering Research Council of Canada.

## Author Contributions Statement

J.S. performed the calculations with help from H.Z.H. and C.S.; J.S. and C.S. wrote the paper with feedback from H.Z.H.; C.S. conceived and supervised the project.

