## Supplementary Information for "Could radical pairs play a role in xenon-induced general anesthesia?"

### Results with two and three xenon atoms in the active site

In contrast to the case of the cryptochrome model, in which the nuclear spins are permitted to take on different isotopic spin values leading to various permutations of nuclear spins, in the experimental case of xenon each isotope was supplied separately by Li *et al.*[1], so here all xenon nuclei in the RPM model were constrained to have the same simultaneous isotopic nuclear spin ($I_1 = I_2 = I_3 \in \{0,\ 1/2,\ 3/2\}$).

In the case of two xenon nuclei occupying the active site the assumption was made that each xenon nucleus couples exclusively to a unique electron in the RP, motivated by the proximity of the xenon nuclei to each phenylalenine residue, and by extension to each radical electron, in the work of Dickinson *et al.* The Hamiltonian describing this case, where each of the two nuclei couples solely to a unique radical electron, can be expressed as

$$\hat{H} = \omega\left(\hat{S}_{Az} + \hat{S}_{Bz}\right) + a_1 \hat{\mathbf{S}}_A \cdot \hat{\mathbf{I}}_1 + a_2 \hat{\mathbf{S}}_B \cdot \hat{\mathbf{I}}_2. \tag{1}$$

The optimized parameter values using the two-xenon occupation state were found to be $a_1' = 0\ \mu$T, $a_2' = 1000\ \mu$T, $B = 50\ \mu$T, $k = 0.2 \times 10^6\ \text{s}^{-1}$, and $r = 1.0 \times 10^6\ \text{s}^{-1}$, resulting in $SR_{1/2} = 0.73$ and $SR_{3/2} = 0.71$. Similar to the single-xenon case, the two-xenon model produced results indicating that the RPM can explain experimental anesthetic potency results most closely in the case that one xenon nucleus couples with its respective radical electron very strongly, while the other xenon nucleus interacts with its corresponding electron relatively weakly, which essentially renders the two cases equivalent; one radical electron couples strongly with one xenon atom, while the other electron only couples weakly. In the two-xenon case, however, the hyperfine parameter space for which *SR* is within uncertainty of *Pot* is relatively large compared with the single-xenon case, as seen in Supplementary Figures S1 and S2. Similar optimized RP lifetime values are seen in this case as in the case of single-xenon occupation, see Supplementary Figs. S3 and S4, but the range of agreeable values extends to somewhat longer RP lifetimes than in the single-xenon case, where both cases are within an order of magnitude of the value suggested by Hore[2] in the cryptochrome context.

The results of the *B*-sensitivity analysis of the two-xenon occupation model show that for external fields with either very small ($< 24\ \mu$T) or extremely large ($> 1102\ \mu$T) magnitudes, *SR* and *Pot* values may diverge, see Supplementary Fig. S5.

Considering the three-xenon occupation state in which three xenon atoms occupy the glycine binding site in the NMDA receptor, based on the geometric modelling done by Armstrong *et al.*[3] it was assumed that radical electron A couples with two xenon nuclei ($I_1$ and $I_2$) and radical electron B couples only to the third xenon nucleus ($I_3$). In this case the Hamiltonian can be modelled as

$$\hat{H} = \omega\left(\hat{S}_{Az} + \hat{S}_{Bz}\right) + a_1 \hat{\mathbf{S}}_A \cdot \hat{\mathbf{I}}_1 + a_2 \hat{\mathbf{S}}_A \cdot \hat{\mathbf{I}}_2 + a_3 \hat{\mathbf{S}}_B \cdot \hat{\mathbf{I}}_3, \tag{2}$$

The results of the three-xenon model suggest that in order to explain the experimental isotopic spin dependence of the anesthetic potency of xenon reported by Li *et al.*[1], the use of the RPM model is idealized in the case that xenon nuclei $I_1$ and $I_3$ have similar hyperfine coupling strengths with radical electrons A and B, respectively, but xenon nucleus $I_2$ would have relatively extremely weak hyperfine coupling with radical electron A. The work of Armstrong *et al.*[3] may provide some insight into this result. In their work, various amino acids in the glycine binding site of NMDA receptors were mutated and the inhibition of the NMDA receptor by xenon was measured. Their results suggest that xenon is more likely to interact with phenylalanine than tryptophan residues, which is consistent with the proposal that a naturally occurring RP may involve the two phenylalanine residues and exclude the tryptophan residue located in the glycine binding site. Their work also predicts that the arrangement of the xenon atoms within the glycine binding site has triangular geometry, as shown in Fig. 1 of the paper, with the xenon atoms positioned such that relative to the two phenylalanine residues there is one xenon nucleus ($I_1$ and $I_3$) in closest proximity to each radical electron, while the third xenon nucleus ($I_2$) is nearest to the tryptophan residue and furthest from the two phenylalanine residues hypothetically forming the RP.

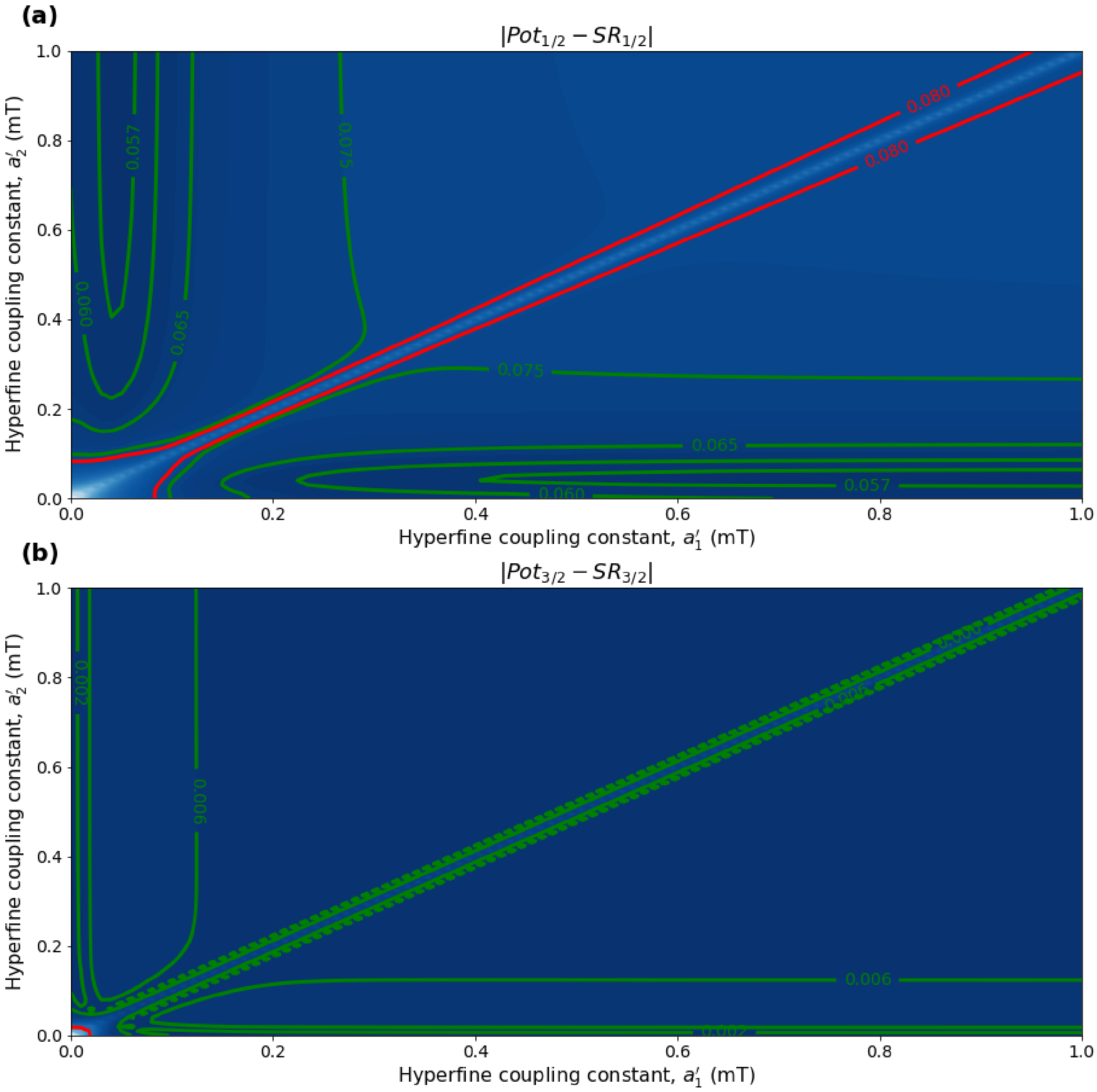


**Supplementary Figure S1.** The dependence of the two-xenon RPM model on changes in the hyperfine coupling constants $a_1'$ and $a_2'$ for $a_1'$, $a_2' \in [0, 1000]$ $\mu$T, using $r = 1.0 \times 10^6$ s$^{-1}$, $B = 50$ $\mu$T, and $\tau = 4.9 \times 10^{-6}$ s. The model can explain the experimentally derived[1] relative anesthetic potency of xenon for values of $r$ and $k$ where $|Pot_{1/2} - SR_{1/2}|$, $|Pot_{3/2} - SR_{3/2}| \leq 0.08$. **(a)** The absolute difference between $Pot_{1/2}$ and $SR_{1/2}$. **(b)** The absolute difference between $Pot_{3/2}$ and $SR_{3/2}$.

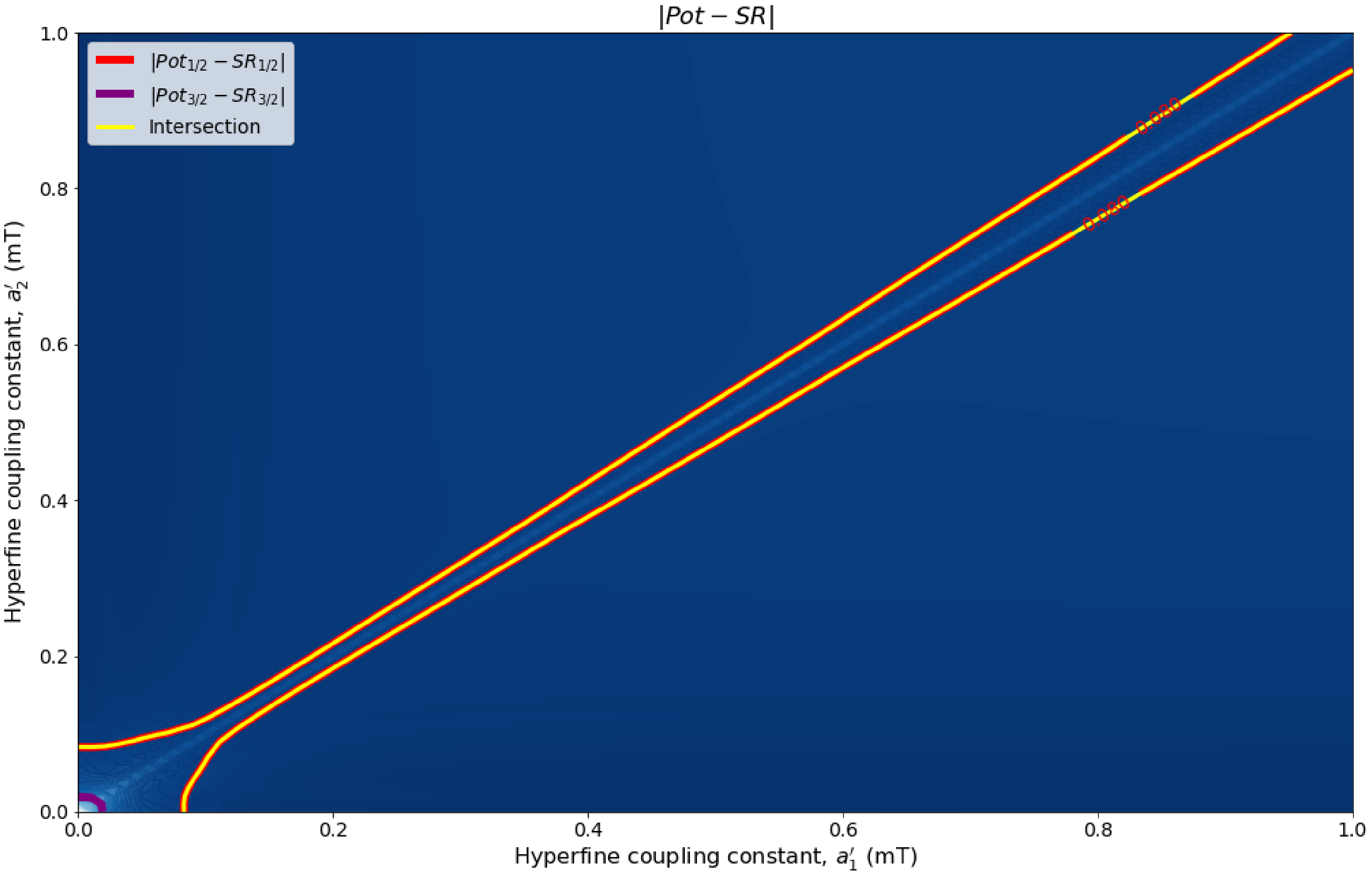


**Supplementary Figure S2.** The dependence of the two-xenon RPM model on changes in the hyperfine coupling constants $a'_1$ and $a'_2$ for $a'_1$, $a'_2 \in [0, 1000]$ $\mu$T, using $r = 1.0 \times 10^6$ s$^{-1}$, $B = 50$ $\mu$T, and $\tau = 4.9 \times 10^{-6}$ s. The model can explain the experimentally derived[1] relative anesthetic potency of xenon where $|Pot_{1/2} - SR_{1/2}| \leq 0.080$ and $|Pot_{3/2} - SR_{3/2}| \leq 0.08$ intersect.

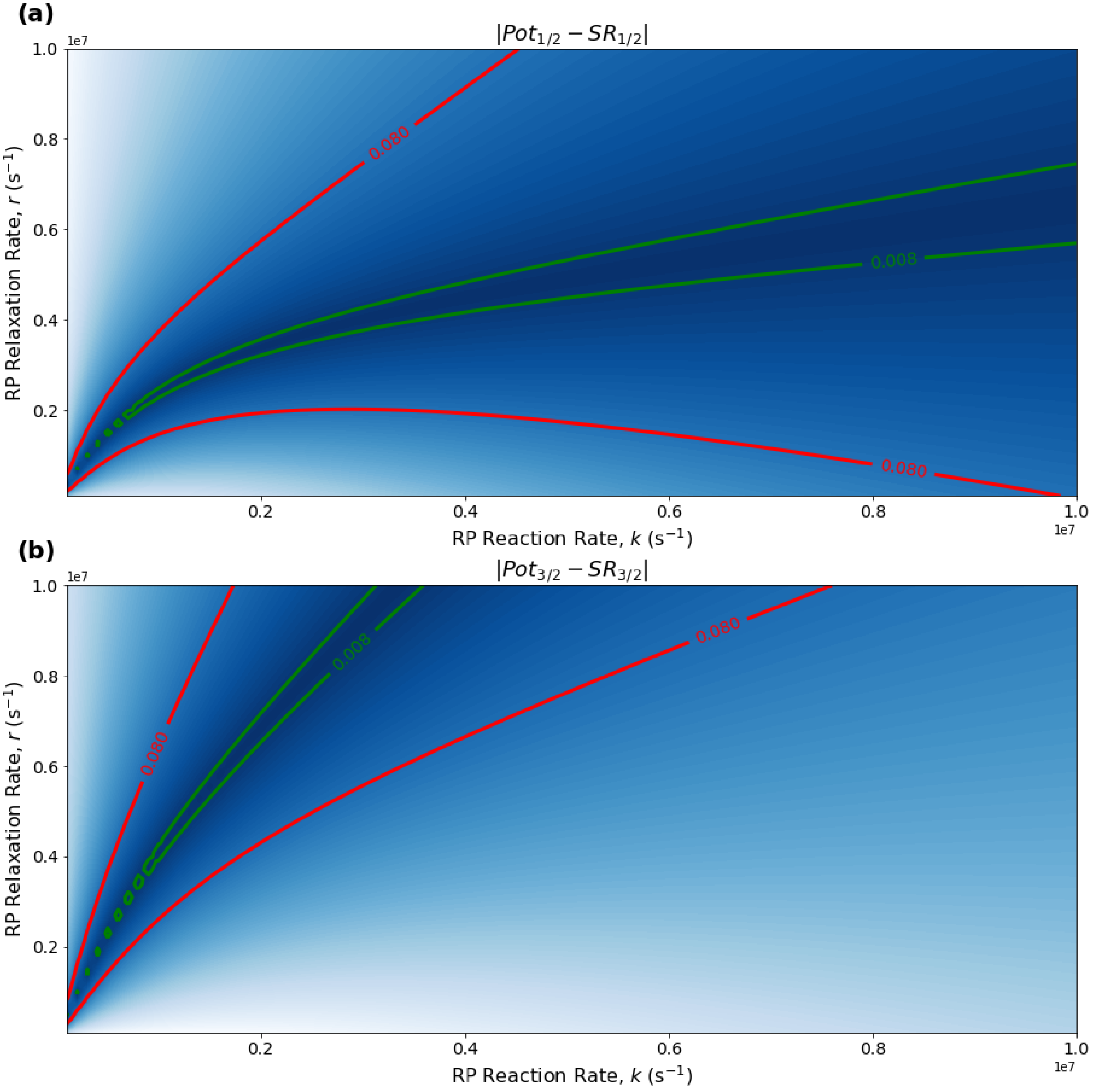


**Supplementary Figure S3.** The dependence of the two-xenon RPM model on the relationship between $r$ and $k$ for $r,\ k \in [1.0\times10^5, 1.0\times10^7]\ \mathrm{s}^{-1}$, using $a_1' = 0\ \mu\mathrm{T}$, $a_2' = 1000\ \mu\mathrm{T}$, and $B = 50\ \mu\mathrm{T}$. The model can explain the experimentally derived[1] relative anesthetic potency of xenon for values of $r$ and $k$ where $|Pot_{1/2} - SR_{1/2}|,\ |Pot_{3/2} - SR_{3/2}| \leq 0.08$. **(a)** The absolute difference between $Pot_{1/2}$ and $SR_{1/2}$. **(b)** The absolute difference between $Pot_{3/2}$ and $SR_{3/2}$.

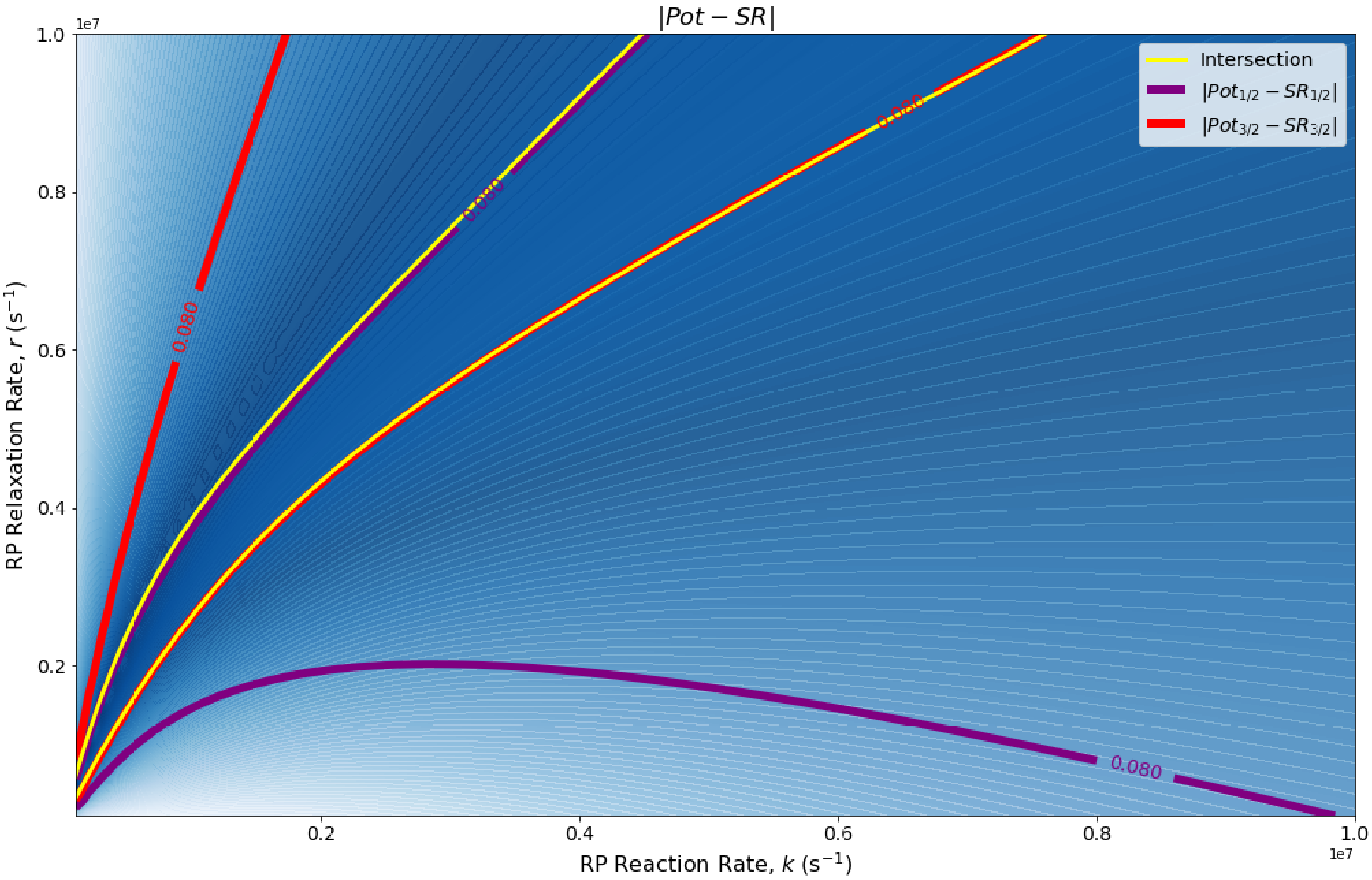


**Supplementary Figure S4.** The dependence of the two-xenon RPM model on the relationship between $r$ and $k$ for $r,\ k \in [1.0\times10^5, 1.0\times10^7]$ $\mathrm{s}^{-1}$, using $a'_1 = 0$ $\mu$T, $a'_2 = 1000$ $\mu$T, and $B = 50$ $\mu$T. The model can explain the experimentally derived[1] relative anesthetic potency of xenon where $|Pot_{1/2} - SR_{1/2}| \leq 0.080$ and $|Pot_{3/2} - SR_{3/2}| \leq 0.08$ intersect.

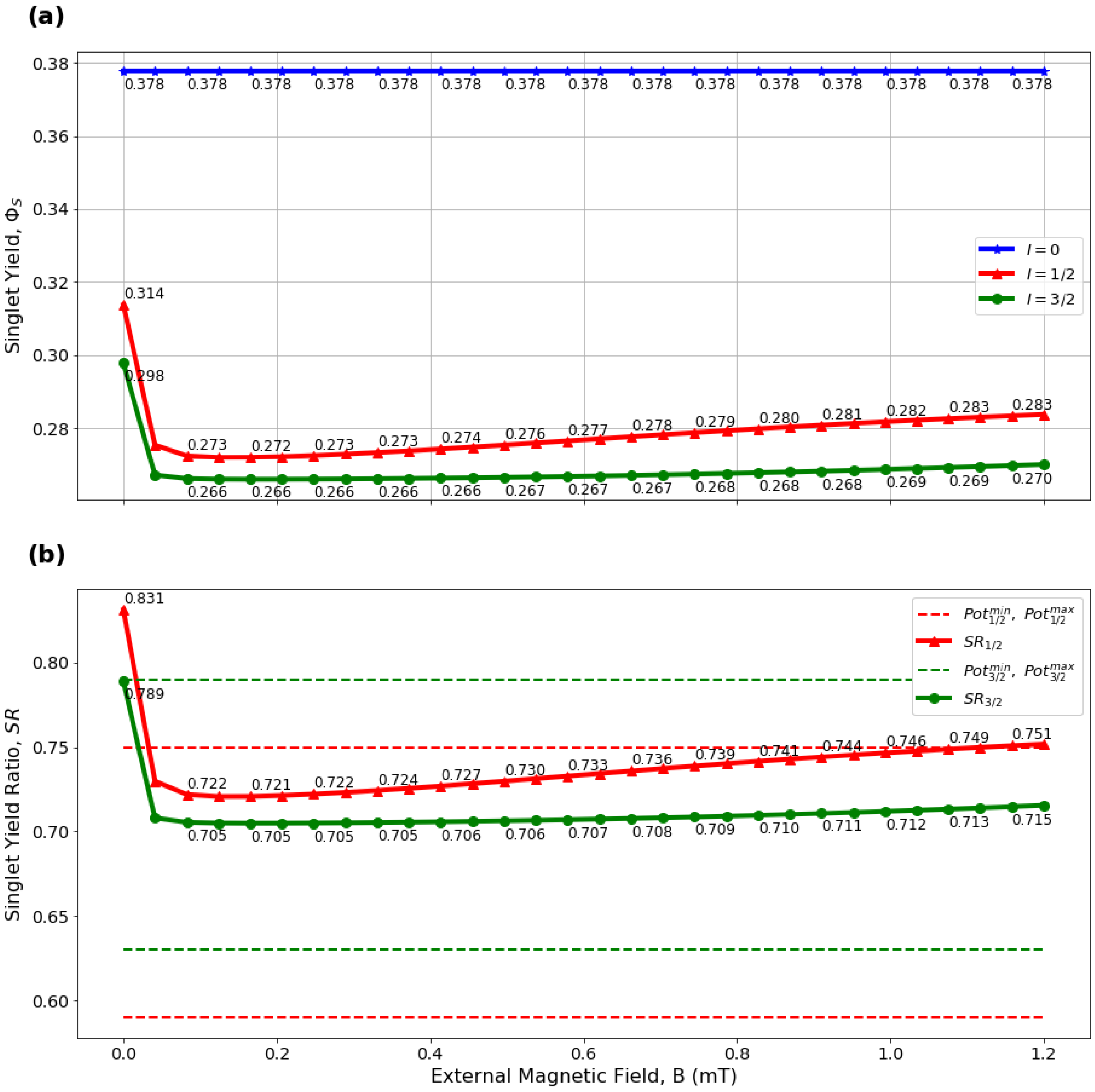


**Supplementary Figure S5.** The external field-dependence of the two-xenon RPM model with $B \in [0,\ 1200]\ \mu$T, $\tau = 4.9 \times 10^{-6}$ s, $r = 1.0 \times 10^6\ \text{s}^{-1}$, and hyperfine constants of $a'_1 = 0\ \mu$T, $a'_2 = 1000\ \mu$T. **(a)** The absolute singlet yield using xenon isotopic nuclear spin values of $I \in \{0,\ 1/2,\ 3/2\}$. **(b)** The relative singlet yield ratios $SR_{1/2}$ and $SR_{3/2}$ with the singlet yield of spin $I = 0$ being normalized to $SR_0 = 1$. Values of $SR$ and $Pot$ agree for $B \in [24,\ 1102]\ \mu$T.
